# Chloroplast genome assemblies and comparative analyses of major *Vaccinium* berry crops

**DOI:** 10.1101/2022.02.23.481500

**Authors:** Annette M. Fahrenkrog, Gabriel Matsumoto, Katalin Toth, Soile Jokipii-Lukkari, Heikki M. Salo, Hely Häggman, Juliana Benevenuto, Patricio Munoz

## Abstract

**Background:** *Vaccinium* is an economically important genus of berry crops in the family Ericaceae. Given the numerous hybridizations and polyploidization events among *Vaccinium* species, the taxonomy of this genus has remained uncertain and the subject of long debate. Therefore, the availability of more genomic resources for *Vaccinium* can provide useful tools for phylogenetic resolution, species identification, authentication of berry food products, and a framework for genetic engineering.

**Results:** In this study, we assembled five *Vaccinium* chloroplast sequences representing the following berry types: northern highbush blueberry (*V. corymbosum*), southern highbush blueberry (*V. corymbosum* hybrids), rabbiteye blueberry (*V. virgatum*), lowbush blueberry (*V. angustifolium*), and bilberry (*V. myrtillus*). Two complete plastid genomes were achieved using long-read PacBio sequencing, while three draft sequences were obtained using short-read Illumina sequencing. Comparative analyses also included other previously available *Vaccinium* chloroplast sequences, especially the commercially important species *V. macrocarpon* (cranberry). The *Vaccinium* chloroplast genomes exhibited a circular quadripartite structure, with an overall highly conserved synteny and sequence identity among them. Despite their high similarity, we identified some polymorphic regions in terms of expansion/contraction of inverted repeats, gene copy number variation, simple sequence repeats, and single nucleotide polymorphisms. Phylogenetic analysis revealed multiple origins of highbush blueberry plastomes, likely due to the hybridization events during northern and southern highbush blueberry domestication.

**Conclusions:** Our results enrich the genomic data availability for new *Vaccinium* species by sequencing and assembling the chloroplast DNA of major economically important berry types. Additional whole plastome analyses including more samples and wild species will be useful to obtain a refined knowledge of the maternal breeding history of blueberries and increase phylogenetic resolution at low taxonomic levels.

## Background

The genus *Vaccinium* L. (family Ericaceae) comprises more than 450 species of wide geographic distribution, occurring mostly in the Northern Hemisphere and in mountainous regions of tropical Asia, Central and South America. With a few exceptions, most of the berry fruits produced by the genus are edible by both birds and mammals [1]. Some species have become economically important crops over the past century, being either bred and cultivated in commercial fields, or harvested from managed wild stands [2]. The major commercial crops are northern highbush blueberries (*V. corymbosum* L.), southern highbush blueberries (*V. corymbosum* L. hybrids), lowbush blueberries (*V. angustifolium* Aiton), rabbiteye blueberries (*V. virgatum* Aiton), bilberries (*V. myrtillus* L.), cranberries (*V. macrocarpon* Aiton), and lingonberries (*V. vitis-idaea* L.). In addition to their pleasant flavors, the nutritional value of these berries has led to a significant increase in consumption and production worldwide. In the United States alone, the wholesale value of the *Vaccinium* berry industry exceeds US$1 billion per year [3].

Given the diversity and complexity of the genus *Vaccinium*, it has been further divided into more than 33 sections or subgenera [4]. The most important *Vaccinium* crop species are found in the sections *Cyanococcus* (blueberries), *Oxycoccus* (cranberry), *Vitis-Idaea* (lingonberry), and *Myrtillus* (bilberry) [5]. However, species and section delimitations have been extensively discussed in the literature, as they do not form monophyletic groups [6, 7]. The taxonomic classification has been difficult to resolve because of considerable phenotypic variability with overlapping morphologies, complex ploidy series (ranging from diploids to hexaploids), and general lack of crossing barriers leading to numerous hybridization events [5]. As a result, some species are burdened with an extensive synonymy according to different authors [1, 8, 9]. Nevertheless, this great diversity and intra-/inter-sectional cross-compatibility have been exploited by breeding programs, allowing for the introduction of useful traits from many species [10–14]. Interspecific hybridizations within the *Vaccinium* section *Cyanococcus*, for example, have played a critical role in the development of low chill southern highbush blueberries through numerous crosses of northern highbush blueberry with warm-adapted Florida native species [10, 15].

A few studies have used molecular data to perform phylogenetic analyses of the genus and relevant sections, including the use of simple sequence repeats [16], chloroplast *matK* and *ndhF* genes and the nuclear ribosomal ITS region [6, 17]. These studies have supported the polyphyletic status of current taxonomic groups and were not able to resolve close relationships. With the decreasing costs of next-generation sequencing, using the whole plastome as a “super-barcode” is becoming a popular strategy for increased resolution at lower plant taxonomic levels [18–20]. Whole chloroplasts can provide more sequence-based polymorphisms, and its genetic properties (i.e., uniparental inheritance, haploid, and non-recombinant nature) can simplify phylogenetic reconstructions when dealing with mixed-ploidy species. However, only a few *Vaccinium* chloroplast genomes have been published so far, with most of these studies reporting only the plastome assembly, without performing comparative analyses [21–27]. Moreover, organellar genomes of horticultural plants are overall underrepresented in databases [28].

By generating additional chloroplast genome sequences for *Vaccinium* species, we aim to provide valuable resources to assist future taxonomic and domestication studies, the development of simple molecular markers for the authentication of berry-based products [29], and a framework for chloroplast biotechnology [30, 31]. Therefore, in this study, we report the assembly of five *Vaccinium* chloroplast sequences representing the following berry types: northern highbush blueberry – NHB (*V. corymbosum*), southern highbush blueberry – SHB (*V. corymbosum* hybrids), rabbiteye blueberry (*V. virgatum*), lowbush blueberry (*V. angustifolium*), and bilberry (*V. myrtillus*). We compared the assemblies in terms of synteny and gene content. We also performed whole plastome phylogenetic analyses including other available *Vaccinium* sequences.

## Results

### Chloroplast genome assembly

The whole genome sequence reads used to assemble the chloroplast DNA (cpDNA) of the five *Vaccinium* species were obtained using two different sequencing platforms: (i) a PacBio long reads approach was used to sequence SHB and rabbiteye, and (ii) an Illumina short reads approach was used for NHB, lowbush, and bilberry.

Complete cpDNA assemblies (sequences without any gaps) were obtained for SHB and rabbiteye using PacBio long reads. A total of 20 contigs (longest contig: 277,507 bp) were assembled for SHB, while the rabbiteye assembly generated two contigs (longest contig: 233,010 bp). Given the length of the complete cpDNA from a related species (cranberry) downloaded from GenBank (∼176 kb) [22], the longest contigs of SHB and rabbiteye were likely to contain the complete cpDNA sequence. When the longest contigs were circularized, redundant sequences from their termini were trimmed. The assemblies were further polished, introducing minor modifications. The final SHB and rabbiteye assemblies were 191,378 and 195,878 bp long, respectively (Table 1).

**Table 1.** Assembly and annotation statistics of the chloroplast genomes from six *Vaccinium* species.

| *cpDNA* | SHB | Rabbiteye | NHB | Lowbush | Bilberry | Cranberry |
| --- | --- | --- | --- | --- | --- | --- |
| **Species** | *V. corymbosum*<br>hybrids | *V. virgatum* | *V.*<br>*corymbosum* | *V.*<br>*angustifolium* | *V. myrtillus* | *V. macrocarpon* |
| **Genotype** | Arcadia | Ochlockonee | Draper | Brunswick | OU-L2 | Stevens |
| **Sequencing** | PacBio/Illumina | PacBio/Illumina<br>a | Illumina | Illumina | Illumina | PacBio |
| **Assembler** | Canu | Canu | Novoplasty | Novoplasty | Spades/CAP3 | Canu |
| **Genome size (bp)** | 191,378 | 195,878 | 186,057 | 182,334 | 191,744 | 176,095 |
| **Number of gaps**<br>**(stretch of Ns)** | 0 | 0 | 2 | 5 | 16 | 0 |
| **LSC size (bp)** | 106,385 | 106,427 | 105,714 | 107,607 | 107,134 | 104,591 |
| **SSC size (bp)** | 3,037 | 3,035 | 3,027 | 2,997 | 3,008 | 3,028 |
| **IR size (bp)** | 40,978 | 43,208 | 38,658 | 35,865 | 40,801 | 34,238 |
| **Total number of**<br>**genes\*** | 112 (136) | 112 (136) | 112 (136) | 112 (139) | 112 (145) | 112 (134) |
| **Protein-coding**<br>**genes\*** | 74 (85) | 74 (85) | 74 (85) | 74 (85) | 74 (89) | 74 (85) |
| **tRNA genes\*** | 34 (43) | 34 (43) | 34 (43) | 34 (46) | 34 (48) | 34 (41) |
| **rRNA genes\*** | 4 (8) | 4 (8) | 4 (8) | 4 (8) | 4 (8) | 4 (8) |
| **GC%** | 36.8 | 36.8 | 36.8 | 36.8 | 36.6 | 36.8 |
| **Accession Number** | XXXX | XXXX | XXXX | XXXX | XXXX | MK715447.1 |
| **Reference** | This work | This work | This work | This work | This work | Diaz-Garcia et al.<br>2019 |

\*Number of unique functional genes. In parentheses: Number of genes including duplicates.

The NHB, lowbush and bilberry cpDNAs were obtained from short reads only, resulting in lower quality assemblies compared to the SHB and rabbiteye cpDNA sequences. The short-read assemblies yielded several contigs and a reference-guided scaffolding was performed to obtain a single pseudomolecule. The polishing procedure and the placement of a consensus inverted repeat into the sequences were collectively able to close some gaps, although a few remained. The final draft cpDNA assemblies had 186,057, 182,334, and 191,744 bp for NHB, lowbush and bilberry, respectively (Table 1).

The final cpDNA sequences obtained showed a quadripartite structure, with a large single copy (LSC) region ranging between 105,715 and 107,608 bp, a pair of inverted repeats (IRA and IRB) ranging from 35,864 to 43,207 bp, and a small single copy (SSC) region ranging from 2,998 to 3,038 bp (Figure 1,3 Table 1, Figure S1). The SSC region was inverted in the cranberry cpDNA compared to the other assemblies (Figure 1, Figure 2).

**Figure 1.**
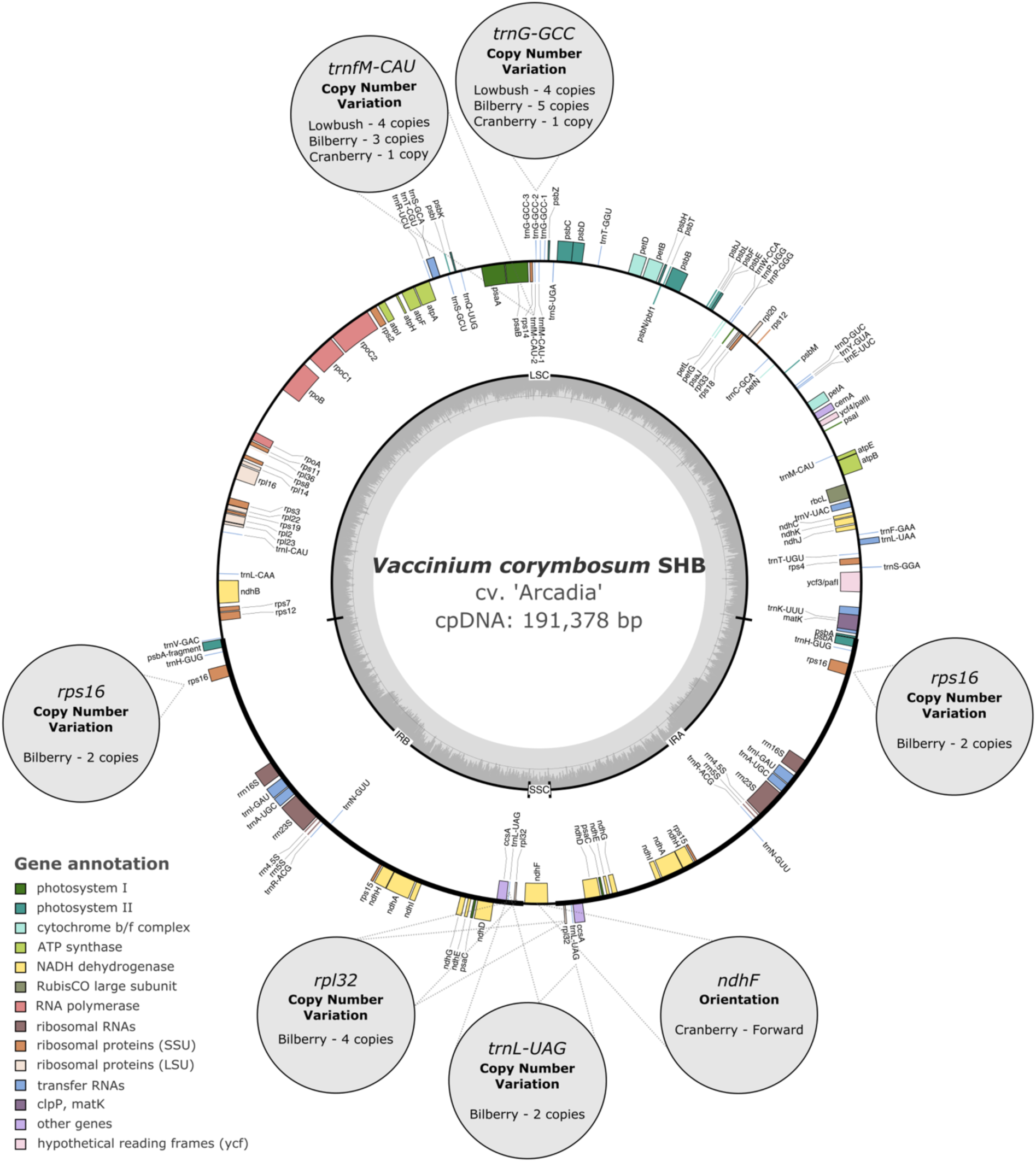
Circular chloroplast genome map of southern highbush blueberry cv. ‘Arcadia’ (*V. corymbosum* hybrids). Outer gray bubbles indicate the variable annotation features among the six *Vaccinium* assemblies. Genes drawn outside and inside the map represent genes transcribed counterclockwise and clockwise, respectively, and the different colors represent their putative functional annotation. The large single copy (LSC), inverted repeats (IRA and IRB), and small single copy (SSC) regions are shown in the black inner circle. The gray inner circle shows GC content.

**Figure 2.**
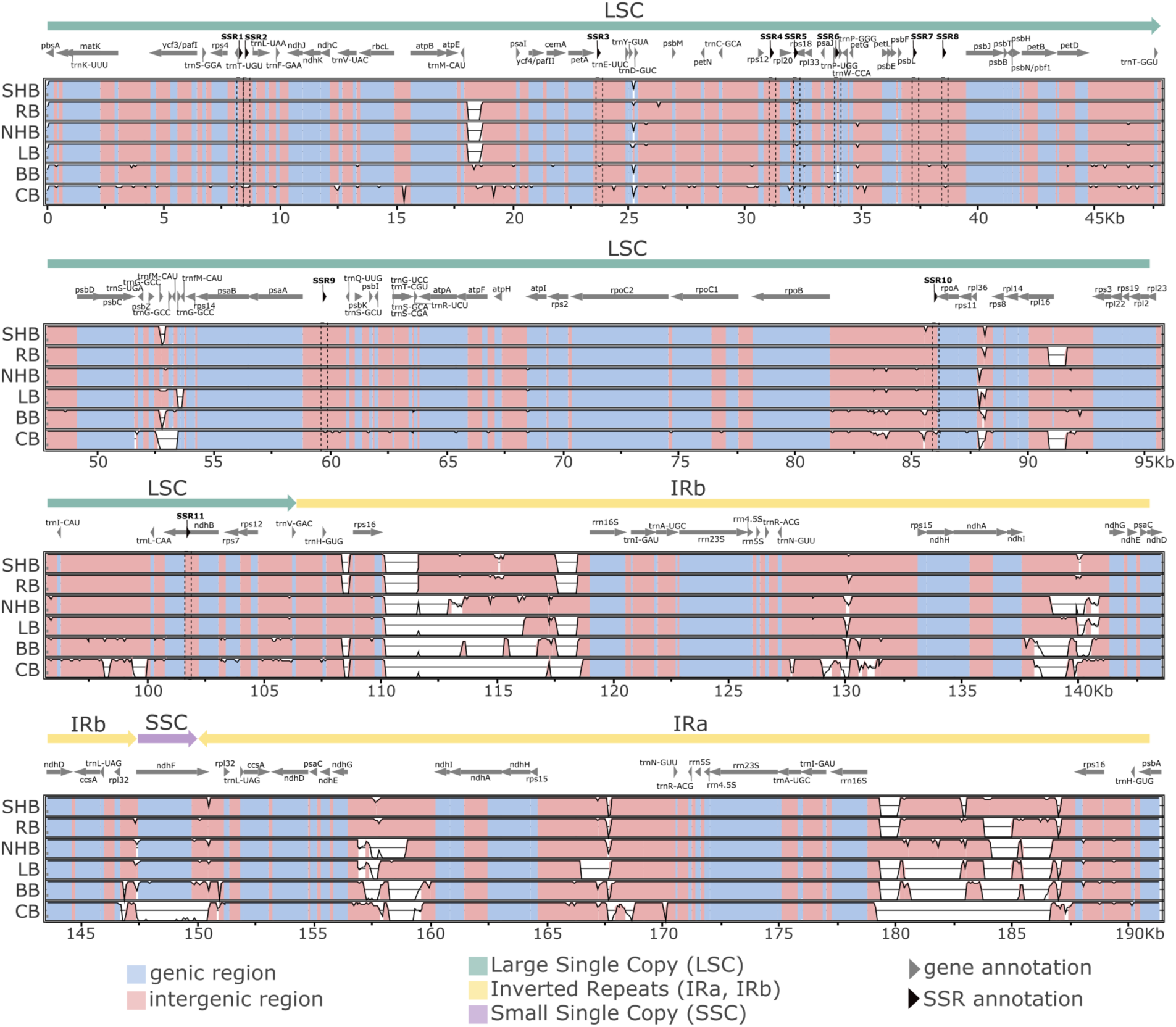
Multiple sequence alignment of *Vaccinium* chloroplast genomes performed with mVISTA. The x-axis represents the coordinates of the southern highbush blueberry chloroplast genome sequence used as reference. The y-axis represents the percentage identity ranging from 50 to 100% for each *Vaccinium* species. Divergent regions due to low sequence similarity or presence of insertions/deletions are shown in white. Polymorphic simple sequence repeat (SSR) regions are highlighted with dashed lines. Abbreviations correspond to the species common names as following: SHB (Southern Highbush Blueberry), RB (Rabbiteye Blueberry), NHB (Northern Highbush Blueberry), LB (Lowbush Blueberry), BB (Bilberry), and CB (Cranberry).

### Gene annotation

The five cpDNA sequences assembled here (SHB, NHB, rabbiteye, lowbush, and bilberry) and the cranberry cpDNA assembly downloaded from GenBank (with minor modifications, see Methods) were annotated for genic features, including ribosomal RNAs (rRNAs), transfer RNAs (tRNAs), and protein-coding genes. For all samples, around 40% of the annotated features had to be manually curated by comparison with annotations available for other plant species (Table S2).

All chloroplast genomes contained the same number of unique putatively functional genes (112), including 74 protein-coding genes, 34 tRNAs, and 4 rRNAs (Table 1 and Table S2). However, the genomes differed in the number of copies present for the genes *rpl32, rps16, trnfM-CAU, trnG-GCC*, and *trnL-UAG* (Figure 1, Table S2). Most of the copy number variation of genes occurred in the draft sequences of lowbush and bilberry. The LSC contains most of the tRNAs (28) and protein-coding genes (63). The IRs contain all four rRNA genes, 11 protein-coding genes and six tRNA genes, which are therefore duplicated in the chloroplast genomes. The SSC contains only one protein-coding gene (*ndhF*), transcribed in the opposite orientation in cranberry when compared to the other genomes (Figure 1).

Nineteen genes contain introns: ten protein-coding genes, and nine tRNA genes. Among those genes, the *rps12* and *psbA* genes had interesting patterns. For the *rps12* gene, the first exon was predicted to be transcribed in the forward direction, while exons 2 and 3 were encoded in the reverse orientation. The *rps12* gene segment containing exon 1 was separated by around 73 kb from the segment containing exons 2 and 3. The *psbA* gene was the only gene spanning the LSC/IR junction, with the starting portion (236 bp) located in the LSC region and the remaining portion (826 bp) located at the end of the IRA. A fragment of the gene is also present in the IRB region, but this partial copy of *psbA* lacks the gene start. The *psbA* gene segments show the same length in all assemblies except in lowbush, where the gene start located in the LSC is 386 bp long due to an insertion.

In addition to putative functional genes, eight gene fragments or pseudogenes were reported by the annotation programs in the six *Vaccinium* assemblies: *accD, clpP, infA, psbG, ycf1, ycf2, ycf15, ycf68* (Table S3). These gene fragments/pseudogenes were removed from the final annotation files and analyses.

### Comparative genomic analysis

The sequence similarity between the six cpDNAs was assessed through multiple sequence alignments, which showed that the *Vaccinium* cpDNAs are highly conserved and syntenic, with most of the variation present in non-coding regions (Figure 2, Figure S2). The main structural differences found were insertions/deletions around the IR borders and the opposite orientation of the SSC in the cranberry cpDNA when compared to the other assemblies. Overall, the cpDNAs showed a sequence identity to the consensus ranging between 82.98% (cranberry) and 91.50% (rabbiteye). The most conserved regions were the LSC (94.16 – 97.23% of identity) and the SSC (91.53 – 98.51% of identity), while the IR was the most divergent region (69.82 – 87.64% of identity) (Table S4).

### Simple sequence repeat analysis

The six *Vaccinium* cpDNA assemblies were screened for the presence of simple sequence repeats (SSRs), identifying between 77 (lowbush) and 109 (rabbiteye) SSRs (Table S5, Table S6). Mononucleotide repeats were most frequent repeat type found (27-41), followed by tetra-(18-28) and dinucleotide repeats (9-31). Trinucleotide repeats were found in lesser numbers (6-8) and pentanucleotide repeats were identified only in bilberry (4) and cranberry (6). Hexanucleotide repeats were more frequent in SHB (13) and rabbiteye (15) than in the other species (3-9), while compound repeats (two repeats separated by a non-repeat sequence) were more frequent in NHB (14) and lowbush (12) than in the remaining species (2-6). Most compound repeats were composed of mononucleotide repeats. In terms of SSR density, the inverted repeats contained ∼0.75 SSRs/kb, twice as many as the single copy regions (∼0.34 SSRs/kb).

Given that they are easier to genotype and thus are potentially useful as molecular markers, we looked for polymorphisms among the six *Vaccinium* plastomes considering SSRs with di-, tri-, tetra-, penta-, and hexanucleotide repeats, excluding mononucleotide and compound repeats. A total of 54 SSR loci were evaluated either because they were detected in all species, or because they were detected in a subset of species but not in others, indicating that they were missed for not meeting the repeat number detection threshold and thus were likely polymorphic. The ClustalW multiple sequence alignment revealed that most SSRs at the IRs were in regions with gaps for some species and were present multiple times in the *Vaccinium* genomes, making them poor candidates for marker development. In contrast, most SSRs located in the LSC mapped to only one region, yielding a list of 11 polymorphic SSRs (Figure 2). Bilberry and cranberry showed greater variation at these loci, while SHB, rabbiteye, NHB and lowbush generally shared the same alleles (Table S7, Figure S3). Therefore, even the combination of these 11 SSRs was not enough to discriminate among closely related species.

### Phylogenetic tree

The whole plastome alignment of 15 species yielded homologous sequence blocks comprising a total of 86,628 bp in length. Most of the sites were conserved across the species, and 8,205 single nucleotide polymorphisms (SNPs) were detected. Out of those, 1,461 were parsimony-informative and 6,744 were singletons (i.e., mutations appearing only once among the sequences).

A maximum likelihood tree was reconstructed to show the phylogenetic relationships among the species (Figure 3A). *Vaccinium* species belonging to different sections were supported in the phylogenetic tree, except for the *Cyanococcus* section which was not monophyletic. The species *V. uliginosum* is classified as in the section *Vaccinium*, however it was placed among the species in section *Cyanococcus*. Within the Cyanococcus section, it is also noteworthy that the SHB cpDNA was more closely related to *V. virgatum* (rabbiteye) than to *V. corymbosum* (NHB), while NHB showed a closer relationship to *V. angustifolium* (lowbush blueberry).

**Figure 3.**
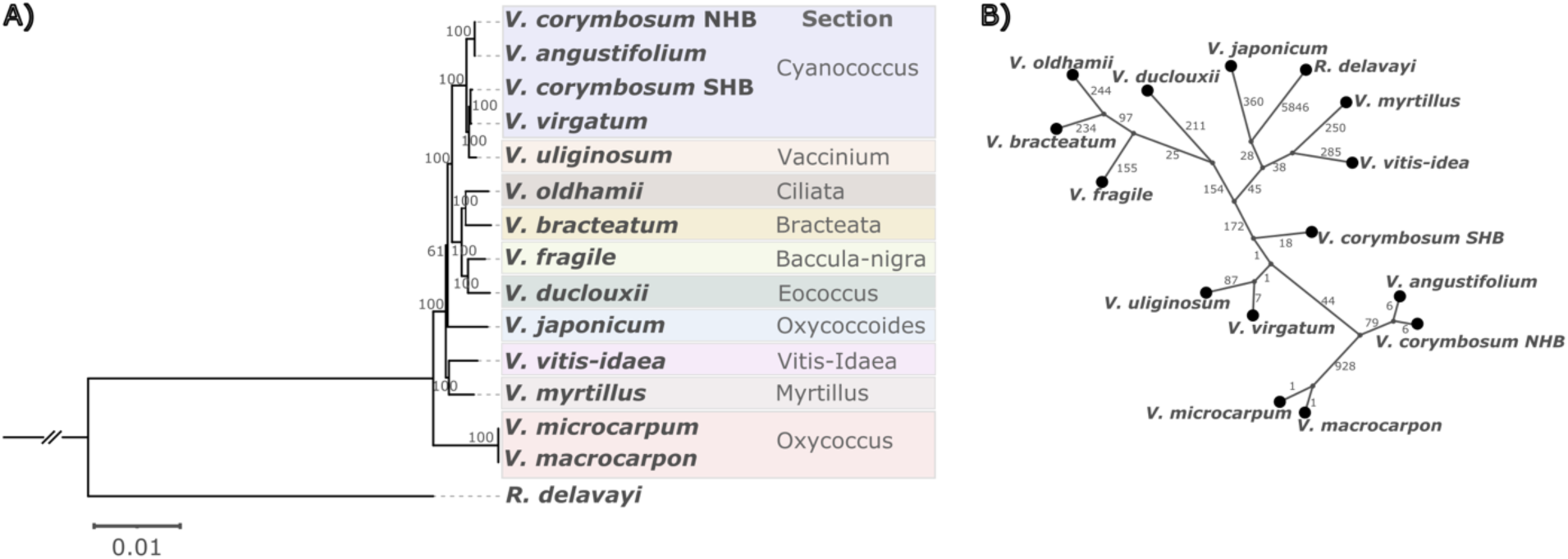
Phylogenetic and haplotype network analyses of whole chloroplast genomes of *Vaccinium* species. **A)** Maximum likelihood phylogenetic tree. Different shades of colors represent different *Vaccinium* sections. Branch labels indicate the bootstrap support values. The scale bar represents nucleotide substitutions per site and double slash-marks indicate out-of-scale. The sequence of *Rhododendron delavayi* was used as an outgroup to root the tree. **B)** Haplotype network showing the mutational steps separating the *s*pecies. Segment length is not proportional to number of mutations.

Despite considering a large chloroplast genomic region, only few mutational steps separated haplotypes of closely related species. For example, 27 polymorphisms differentiated SHB from rabbiteye, 12 between NHB and lowbush, and two between cranberry and its wild relative *V. microcarpum* (Figure 3B). Some allelic variants at the tips of the network were species-specific and could serve as potential molecular markers.

## Discussion

Since the first *Vaccinium* chloroplast DNA sequence was published in 2013 [21], next-generation sequencing technologies have enabled the assembly of plastomes for additional species in this genus, making nine *Vaccinium* cpDNAs available to date [21–27]. Here, we performed *de novo* assembly of the plastomes of five additional *Vaccinium* species, including the four most important cultivated blueberry types and bilberry.

The highest quality complete plastome assemblies were obtained for SHB and rabbiteye, which were sequenced using long reads from the PacBio platform. The availability of long reads has allowed assembly of the entire cpDNA as a single contig, similar to the assembly done for cranberry using the same technology [22]. Although the remaining species were sequenced with Illumina short reads and the assemblies were split into more than one contig, the use of a reference cpDNA to order the contigs was able to generate draft plastomes for NHB, lowbush and bilberry containing only a few gaps in their sequences.

All six *Vaccinium* cpDNA sequences compared here showed the typical circular quadripartite structure for angiosperms, including the two copies of inverted repeats separating the large and the small single copy regions [32]. The length of the *Vaccinium* cpDNA assemblies was also within the range reported for plant species (107–218 kb) [20]. However, a drastic reduction in the SSC region was observed among the *Vaccinium* assemblies (∼3 kb) compared to most angiosperms (16–27 kb), which has also been reported for other members of the *Ericaceae* family [33–35].

Most angiosperm chloroplast genomes contain 110–130 distinct genes, approximately 80 genes coding for proteins and other genes coding for 4 rRNAs and 30 tRNAs. For the six *Vaccinium* species analyzed in this study, a total of 112 distinct genes were annotated (74 protein-coding, 34 tRNA and 4 rRNA genes). A recent study comparing the plastomes of five other *Vaccinium* species showed differences in gene content among them [36]. This differs from our findings, where even the most distant taxon in the phylogenetic tree (cranberry) carries the same genes as the other *Vaccinium* species sequenced herein. This discrepancy could be due to software mispredictions. In our study, we observed that using more than one gene prediction software and performing manual curation were important steps for the proper identification of genes in chloroplast genomes. For example, in our work, we identified four tRNA genes (*trnfM-CAU, trnG-GCC, trnS-CGA, trnS-GCA*) not previously reported in the cranberry plastome, and five putatively functional genes (*atpF, ccsA, ndhG, ndhK, rps16*) that were previously considered pseudogenes [21, 22]. Instead of a difference in the absolute number of distinct genes, we found copy number variation for five genes. However, these copy number variations warrant further validation, since most of them were identified in the lower-quality assemblies of lowbush and bilberry. Eight gene fragments or putative pseudogenes were identified here, including the *accD, clpP* and *infA* genes, which have been previously reported as pseudogenes in cranberry but as functional in other members of the *Ericaceae* family[34].

Besides the gene content, comparative genomics analyses among the six *Vaccinium* species also revealed high similarity in terms of sequence identity and synteny. Overall, sequence identity was higher in coding than in non-coding regions. One synteny difference identified was the opposite orientation of the SSC in cranberry when compared to the other assemblies. However, it has been shown in other plant species that both SSC orientations can be present simultaneously within the same individual due to chloroplast heteroplasmy [37, 38]. Therefore, at this point, we cannot consider the SSC orientation a consistent rearrangement in cranberry. Another structural difference was found in a non-coding region close to the IR/LSC boundaries, where the cranberry cpDNA (shortest plastome) shows a missing fragment of approximately 8 kb when compared to rabbiteye (longest plastome). This difference between assemblies is reflected in the lower sequence identity shown within the IRs. Greater sequence divergence within the IRs was also reported in a previous comparison between five other *Vaccinium* species [39]. Indeed, expansion/contraction of the IRs is one of the major causes for plastome size differences between plant species [32].

Comparison of the abundance of different SSR repeat units showed that mononucleotide repeats were the most frequent repeat type. Also, most compound repeats were composed of mononucleotide repeats. Mononucleotide repeats have been shown to be the most abundant and variable class in other plant species [40, 41]; however, their use as molecular markers have been limited given their lower reliability and difficulty to genotype [42]. Therefore, we searched for variability only at orthologous SSRs with longer repeat units and located at single copy regions, identifying 11 polymorphic SSRs. However, these SSRs were unable to distinguish close *Vaccinium* species. Their use for intraspecific variability also needs further investigation. In contrast, some species-specific SNPs were detected throughout the whole cpDNA alignment. They could serve as potential molecular markers, especially for berry food product authentication, as we included the major economically important *Vaccinium* species in the analyses.

In this study, the whole cpDNA phylogenetic analysis was able to distinguish the species of the genus *Vaccinium* and most of the sections were monophyletic. However, the phylogenetic relationships in the *Vaccinium* genus are complex, especially when analyzing domesticated genotypes. Incongruences between the chloroplast phylogeny and previous nuclear phylogenies can be pointed out for cultivated blueberries. The chloroplast genomes of the NHB and SHB genotypes used herein have different origins, with NHB being more closely related to lowbush, and SHB to rabbiteye. In the phylogenetic trees derived from nuclear genome-wide SNPs [43, 44] and SSRs [45], SHB and NHB genotypes were intertwined and more closely related to each other than to lowbush or rabbiteye. Given the primary contribution of *V. corymbosum* to the genetic background of both NHB and SHB [46], it is expected that the nuclear genome would reflect the described pattern. On the other hand, the cpDNA will trace back the maternal line inheritance. As interspecific hybridizations have been extensively used in blueberry breeding programs, lowbush and rabbiteye lineages are present in the NHB and SHB genetic background as secondary gene pools. *V. angustifolium* has been used since the beginning of highbush blueberry domestication [47], with genotypes such as ‘Russell’ and ‘North Sedgewick’ being widely used in crosses. During the development of SHB, several rabbiteye genotypes were used as parents to reduce the chilling requirement of NHB [11, 48, 49]. Therefore, different cultivars of NHB and SHB will likely show different plastome clustering patterns based on the maternal pedigree. Expanding this study to additional wild *Vaccinium* species and multiple individuals would help to clarify their hybridization history and avoid unclear clustering of single accessions [50]. The chloroplast phylogeny also highlights the placement of *V. uliginosum* section *Vaccinium* among species in the section *Cyanococcus*. Intersectional crosses have generally proved difficult to perform, yielding mostly sterile hybrids [51]. However, successful crosses between *V. uliginosum* and *V. corymbosum* were reported to produce meiotically regular and fruitful hybrids [52], which reinforces their closer phylogenetic proximity.

## Conclusions

In this study, the chloroplast genomes of five economically important *Vaccinium* species were assembled: northern highbush blueberry, southern highbush blueberry, rabbiteye blueberry, lowbush blueberry, and bilberry. We also performed manual curation of gene annotations and comparative analyses of these genomes, including the previously available cranberry plastome sequence. The *Vaccinium* chloroplast genomes were highly conserved in terms of structure and sequence, with some variability found mostly in non-coding regions and at the IR/LSC boundaries. Copy number variation of genes requires further investigation as they could be a result of assembly artifacts in draft genomes. Species-specific allelic variants were found for SNPs, but not for SSRs. The phylogenetic tree based on whole cpDNA alignment showed the presence of distinct maternal genomes in highbush blueberries, highlighting the independent evolution of cytoplasmic and nuclear genomes. In addition, chloroplast phylogenetic analyses did not support the monophyly of the *Cyanococcus* section. The availability of more chloroplast genomes from *Vaccinium* species will provide a valuable resource for future comparative studies and phylogenetic resolution of the genus, and for reconstructing the domestication history of cultivated berry crops.

## Methods

### Plant material

The plant material used to generate the DNA sequences for the assembly of the chloroplast DNAs (cpDNAs) included the *V. corymbosum* hybrid cv. ‘Arcadia’ (Southern Highbush Blueberry - SHB), *V. virgatum* cv. ‘Ochlockonee’ (Rabbiteye Blueberry - RB), and *V. angustifolium* cv. ‘Brunswick’ (Lowbush Blueberry - LB) obtained from commercial nurseries and maintained at the University of Florida, FL, USA. The plant material for the *V. myrtillus* genotype ‘OU-L2’ (Bilberry -BB) was collected from the coniferous forest in the municipality of Oulu, Finland (64°59’08.1”N 25°54’12.0”E) and maintained at the University of Oulu. No special permission was required for sampling the bilberry individual at this location. The genomic sequences used to assemble the cpDNA of the *V. corymbosum* cv. ‘Draper’ (Northern Highbush Blueberry - NHB) were downloaded from the Sequence Read Archive (SRA) under the BioProject PRJNA494180 [53]. The cpDNA sequence assembly for *V. macrocarpon* cv. ‘Stevens’ (Cranberry - CB) was downloaded from GenBank (MK715447.1) [22].

### DNA extraction and sequencing

The high molecular weight DNA extraction from young leaf tissue and the PacBio long read sequencing for the SHB and rabbiteye samples were carried out at the Arizona Genomics Institute, University of Arizona (Tucson, AZ, USA). Briefly, the high molecular weight DNA was extracted using a modified CTAB method and sheared to mode size of approximately 40 kb using G-Tube. PacBio sequencing libraries were constructed using the Express v2 kit (Pacific Biosciences). Template molecules were size selected on BluePippin for either 35 kb and larger (U1) or 20 kb and larger (S1) methods (Sage Sciences). Sequencing was performed on PacBio Sequel II, in CLR mode with a loading concentration of 50 pmol or larger. PacBio consumables used were PacBio SeqII 1.0 chemistry, 8Mv1 cells and 15 hr run time.

Short-read Illumina whole genome sequencing was obtained for the SHB, rabbiteye, lowbush and bilberry samples by extracting genomic DNA from leaf tissue using the CTAB method. DNA library preparation and sequencing were carried out at GENEWIZ LCC. (South Plainfield, NJ, USA). Paired end libraries (2×150 bp) were sequenced on an Illumina HiSeq4000 instrument. For bilberry, Illumina paired end library preparation and sequencing was conducted at Sequentia Biotech SL (Barcelona, Spain), using NovaSeq 6000 instrument (2×150 bp). The mean insert size for SHB, rabbiteye, lowbush and bilberry was 325 bp, while the Illumina sequencing data downloaded for NHB included libraries with five different insert sizes: 470 bp, 800 bp, 4,000 bp, 7,000 bp and 10,000 bp (Table S1).

### Long-read assembly and polishing

PacBio long reads from SHB and rabbiteye were aligned to the reference cranberry cpDNA sequence using BLASR v.20130815 with parameters “--placeGapConsistently, --hitPolicy randombest, --bestn 1, --minMatch 15, and --minAlnLength 500” [54]. The aligned sequence was converted into FASTQ format using the function “bamtofastq” from bedtools v2.29.2 software [55]. The retrieved reads were assembled with Canu v1.9 using the parameters “minReadLength=1000, minOverlapLength=500, genomeSize=200k, correctedErrorRate=0.030, and corOutCoverage=40” [56]. The longest contig generated by Canu was circularized using Circlator v.1.5.5 [57] and polished with the Arrow algorithm implemented in the GCpp v1.9.0 software [58]. Five total rounds of polishing with Arrow were performed for SHB and rabbiteye before moving to a second polishing method. The second polishing step was performed with the software Pilon v1.22 [59] using Illumina short reads and default parameters until no more changes were introduced into the sequence (for up to five successive rounds).

### Short-read assembly and polishing

For NHB and lowbush samples, short Illumina read assemblies were performed with the NovoPlasty v3.8.3 software with default parameters [60]. The resulting scaffolds were aligned to the SHB cpDNA assembly obtained previously with long-read data, using the “nucmer” tool available in Mummer v4.0 [61]. The pairwise alignments were visualized using the Mummer tools “show-coords” and “mummerplot” and the individual NovoPlasty scaffolds were ordered and merged into one pseudo-molecule for each sample according to their placement along the reference SHB cpDNA assembly. A stretch of Ns was inserted at the junction sites between concatenated scaffolds.

A similar strategy was used for the bilberry assembly. Raw short Illumina reads were aligned to the cpDNA sequence of *Vaccinium oldhamii* (GenBank accession: NC_042713.1) [24]. The mapped reads were then extracted, and de novo assembled with Spades 3.15.3 [62] and with CAP3 v.20120705 [63]. The two assemblies were then aligned to the reference *V. oldhamii* genome, the scaffolds were ordered and then merged into one pseudo-molecule.

The NHB, lowbush, and bilberry assemblies were polished using Pilon v1.22 as described above. The NHB and lowbush assemblies were polished multiple times, until no further changes were introduced into the sequences (i.e., four and three rounds, respectively). The bilberry assembly was subjected to only one round of polishing, because additional rounds inserted sequences into multiple sites, generating tandem repeats.

To obtain a more continuous sequence in the inverted repeat (IR) regions, for each species the sequences of IRA and IRB were aligned, and the consensus sequence was inserted back into the cpDNA assembly to replace the original IR sequences. Finally, when comparing the IR sequence length in the cranberry assembly downloaded from GenBank, we noticed that two bases were absent from one of the IRs. These nucleotides were inserted into the IR where they were missing, resulting in both IRs having the same length and sequence in the cranberry cpDNA.

### Gene and SSR annotation

The cpDNA sequences were annotated to predict gene content and position. Two online tools were employed: (i) GeSeq v2.03 by setting parameters “protein search identity= 70; rRNA, tRNA, DNA search identity =85; and selecting the 3^rd^ party tRNA annotators ARAGORN v1.2.38 and tRNAscan-SE v2.0.5” [64]; and (ii) CpGAVAS with default parameters [65]. The annotations obtained with both methods were not consistent for many genes. Discordant annotations were manually curated by comparing the software outputs with gene models available for other species in the CpGDB database [66]. The gene sequences predicted for the *Vaccinium* species were compared to the sequence reported in the CpGDB for *Vaccinium* and other model species, including *V. macrocarpon, Vaccinium oldhamii, Arabidopsis thaliana, Brassica napus, Amborella trichopoda*, and *Populus trichocarpa*. Manual curation of gene features was performed for the five *Vaccinium* cpDNAs assembled in this study and for the cranberry cpDNA.

Considering the potential importance of Simple Sequence Repeats (SSRs) in generating genomic diversity, the cpDNA assemblies were annotated using the MISA-web v2.1 software [67]. The minimum number of repetitions was set at ten for mononucleotide repeats, five for dinucleotide repeats, four for trinucleotide repeats, and three for tetra, penta-, and hexa-nucleotide repeats. Orthologous SSRs were inspected for polymorphisms by looking at the multiple sequence alignments (see below).

### Comparative analyses

To investigate the genome structure of the cpDNAs, circular maps were drawn using OGDRAW v1.3.1 [68]. The cpDNA assemblies were compared by conducting multiple sequence alignments using mVISTA with the LAGAN mode [69] and with the EMMA tool in the EMBOSS v6.5.7 software [70] using the ClustalW v.2.1 aligner [71]. The online tool Multiple Sequence Alignment Viewer v1.21.0 [72] was used to visualize alignments generated with EMMA and to estimate the percentage of identity between sequences. Prior to conducting these multiple sequence alignments, the cpDNA sequences were modified to break their circular DNA molecules at the same site as the cranberry cpDNA to ensure that the alignments would start at the same position.

### Phylogenetic analysis

To infer the phylogenetic relationships among our sequences and other available chloroplast genomes from *Vaccinium* species, we downloaded the GenBank sequences of *V. oldhami* (NC_042713.1), *V. bracteatum* (LC521967.1), *V. duclouxii* (MK816300.1), *V. fragile* (MK816301.1), *V. uliginosum* (LC521968.1), *V. japonicum* (MW006668.1), *V. microcarpum* (MK715444.1), and *V. vitis-idaea* (LC521969.1). The sequence of *Rhododendron delavayi* (MN413198.1) was used as an outgroup to root the tree. Given that we did not perform manual gene curation on these other assemblies, we used a whole chloroplast genome alignment approach to avoid variations due to misprediction. For this, all assemblies were reordered to start with the *rbcL* gene sequence using Circlator v.1.5.5 [57]. We used the HomBlocks pipeline to align the whole cpDNA genomes and determine locally collinear blocks among them [73]. The final concatenated alignment length was 86,628 bp divided into eight blocks. The best substitution model based on the Bayesian Information Criterion (BIC) for all eight blocks was “TVM+G”, computed using PartitionFinder v.2.1.1.0 [74]. The concatenated alignment and the “TVM+G” model were then used to reconstruct a maximum likelihood phylogenetic tree using IQ-TREE v.2.1.0 [75] with 1000 ultrafast bootstraps [76]. The resulting tree was visualized with iTOL v.6 [77].

To visualize the mutational steps differentiating the *Vaccinium* species and identify species-specific markers, the HomBlocks alignment was also used for haplotype network reconstruction using PopART [78] with TCS method [79].

## Additional files

### Additional file 1: PDF

Title: Supplementary Figures

**Figure S1**. Circular chloroplast genome maps of six cultivated *Vaccinium* species.

**Figure S2**. ClustalW multiple sequence alignment of the complete plastomes of six cultivated *Vaccinium* species.

**Figure S3**. Sequence variability of 11 SSRs identified in six *Vaccinium* species.

### Additional file 2: xlsx

Title: Supplementary Tables

**TableS1 -** Sequencing data downloaded for NHB cv. ‘Draper’

**TableS2-** Functional genes annotated and curated

**TableS3 -** Putative pseudogenes/gene fragments

**TableS4 -** Percentage of identity between Vaccinium plastomes

**TableS5 -** Simple sequence repeats (SSRs) identified in the six Vaccinium plastomes

**TableS6 -** Summary of SSRs

**TableS7 -** SSR loci showing variation among species

## List of abbreviations

SHB: Southern Highbush Blueberry
NHB: Northern Highbush Blueberry
RB: Rabbiteye Blueberry
LB: Lowbush Blueberry
BB: Bilberry
CB: Cranberry
cpDNA: chloroplast DNA
SSR: Simple Sequence Repeat
SNP: Single Nucleotide Polymorphism
ITS: Internal Transcribed Spacer
IRs: Inverted repeats
LSC: Large Single Copy
SSC: Small Single Copy
rRNA: ribosomal RNA
tRNA: transfer RNAs

## Declarations

## Availability of data and materials

The complete chloroplast genomes and annotations were submitted and will be available in the NCBI database. Accession numbers will be added here and at Table 1 upon acceptance.

## Contributions

PM, JB, and HH conceived and supervised the study. JB, KT, SJL, and HMS collected the plant material and performed DNA extraction for sequencing. AMF, GM, and JB performed the analyses and interpreted the data. AFM and JB wrote the manuscript. All authors read, revised, and approved the final manuscript.

## Ethics declarations

### Ethics approval and consent to participate

Not applicable.

### Consent for publication

Not applicable.

## References

1. vander Kloet SP. The genus Vaccinium in North America. Research Branch Agriculture Canada Publ. 1988.

2. Ballington JR. Collection, utilization, and preservation of genetic resources in Vaccinium. HortScience. 2001; 36.

3. The Vaccinium Coordinated Agricultural Project (VacCAP). https://www.vacciniumcap.org/. Accessed 21 Feb 2022.

4. Sleumer H. Vaccinioidee-Studien. Botanische Jahrbücher. 1941; 71.

5. Hancock JF, Lyrene P, Finn CE, Vorsa N, Lobos GA. Blueberries and cranberries. In Hancock JF editor. Temperate fruit crop breeding. Dordrecht: Springer; 2008.

6. Kron KA, Powell EA, Luteyn JL. Phylogenetic relationships within the blueberry tribe (Vaccinieae, Ericaceae) based on sequence data from matK and nuclear ribosomal ITS regions, with comments on the placement of Satyria. American Journal of Botany. 2002;89.

7. vander Kloet SP. Vaccinia gloriosa. In: Small Fruits Review. 2004;3.

8. Camp WH. The North American blueberries with notes on other groups of Vacciniaceae. Brittonia. 1945;5.

9. Weakley AS. Flora of the Southern and Mid-Atlantic States May 2015. https://ncbg.unc.edu/research/unc-herbarium/floras/. 2015; Accessed 21 Feb 2022.

10. Sharpe RH, Darrow GM. Breeding blueberries for the Florida climate. Proceedings of the Florida State Horticultural Society. 1959;72.

11. Darrow GM, Dermen H, Scott DH. A tetraploid blueberry: From a Cross of Diploid and Hexaploid Species. Journal of Heredity. 1949;40.

12. Draper AD. Draper, A. D. “Tetraploid hybrids from crosses of diploid, tetraploid, and hexaploid Vaccinium species. Acta Horticulturae. 1977; 61.

13. Ballington JR. The role of interspecific hybridization in blueberry improvement. Acta Horticulturae. 2009;810.

14. Vorsa N, Johnson-Cicalese J, Polashock J. A blueberry by cranberry hybrid derived from a Vaccinium darrowii x (V. macrocarpon x V. oxycoccos) intersectional cross. Acta Horticulturae. 2009;810.

15. Lyrene PM. Value of various taxa in breeding tetraploid blueberries in Florida. Euphytica. 1997;94.

16. Schlautman B, Covarrubias-Pazaran G, Fajardo D, Steffan S, Zalapa J. Discriminating power of microsatellites in cranberry organelles for taxonomic studies in Vaccinium and Ericaceae. Genetic Resources and Crop Evolution. 2017;64.

17. Powell EA, Kron KA, Liston A. Hawaiian blueberries and their relatives - A phylogenetic analysis of Vaccinium sections Macropelma, Myrtillus, and Hemimyrtillus (Ericaceae). Systematic Botany. 2002;27.

18. Parks M, Cronn R, Liston A. Increasing phylogenetic resolution at low taxonomic levels using massively parallel sequencing of chloroplast genomes. BMC Biology. 2009;7.

19. Ma PF, Zhang YX, Zeng CX, Guo ZH, Li DZ. Chloroplast phylogenomic analyses resolve deep-level relationships of an intractable bamboo tribe Arundinarieae (Poaceae). Systematic Biology. 2014;63.

20. Daniell H, Lin CS, Yu M, Chang WJ. Chloroplast genomes: Diversity, evolution, and applications in genetic engineering. Genome Biology. 2016;17.

21. Fajardo D, Senalik D, Ames M, Zhu H, Steffan SA, Harbut R, et al. Complete plastid genome sequence of Vaccinium macrocarpon: Structure, gene content, and rearrangements revealed by next generation sequencing. Tree Genetics and Genomes. 2013;9.

22. Diaz-Garcia L, Rodriguez-Bonilla L, Smith T, Zalapa J. Pacbio sequencing reveals identical organelle genomes between american cranberry (Vaccinium macrocarpon ait.) and a wild relative. Genes. 2019;10.

23. Kim Y, Shin J, Oh DR, Kim DW, Lee HS, Choi C. Complete chloroplast genome sequences of Vaccinium bracteatum Thunb., V. vitis-idaea L., and V. uliginosum L. (Ericaceae). Mitochondrial DNA Part B: Resources. 2020;5.

24. Kim SC, Baek SH, Lee JW, Hyun HJ. Complete chloroplast genome of Vaccinium oldhamii and phylogenetic analysis. Mitochondrial DNA Part B: Resources. 2019;4.

25. Chen X, Liu Q, Guo W, Wei H, Wang J, Zhu D, et al. The complete chloroplast genome of Vaccinium duclouxii, an endemic species in China. Mitochondrial DNA Part B: Resources. 2019;4.

26. Guo W, Luo L, Huang Y, Li G, Wang X, Cheng T, et al. The complete chloroplast genome of Vaccinium fragile (Vacciniaceae), a shrub endemic to China. Mitochondrial DNA Part B: Resources. 2019;4.

27. Cho WB, Han EK, Choi IS, Son DC, Chung GY, Lee JH. The complete plastid genome sequence of Vaccinium japonicum (Ericales: Ericaceae), a deciduous broad-leaved shrub endemic to East Asia. Mitochondrial DNA Part B: Resources. 2021;6.

28. Wang X, Cheng F, Rohlsen D, Bi C, Wang C, Xu Y, et al. Organellar genome assembly methods and comparative analysis of horticultural plants. Horticulture Research. 2018;5.

29. Salo HM, Nguyen N, Alakärppä E, Klavins L, Hykkerud AL, Karppinen K, et al. Authentication of berries and berry-based food products. Comprehensive Reviews in Food Science and Food Safety. 2021;20.

30. Jin S, Daniell H. The Engineered Chloroplast Genome Just Got Smarter. Trends in Plant Science. 2015;20.

31. De-la-Peña C, León P, Sharkey TD. Editorial: Chloroplast Biotechnology for Crop Improvement. Frontiers in Plant Science. 2022;13.

32. Jansen RK, Ruhlman TA. Plastid Genomes of Seed Plants. In Ralph Bock and Volker Knoop Editors. Genomics of chloroplasts and mitochondria. Dordrecht, Springer. 2012. p103–126.

33. Martínez-Alberola F, del Campo EM, Lázaro-Gimeno D, Mezquita-Claramonte S, Molins A, Mateu-Andrés I, et al. Balanced gene losses, duplications and intensive rearrangements led to an unusual regularly sized genome in Arbutus unedo chloroplasts. PLoS ONE. 2013;8.

34. Logacheva MD, Schelkunov MI, Shtratnikova VY, Matveeva M v., Penin AA. Comparative analysis of plastid genomes of non-photosynthetic Ericaceae and their photosynthetic relatives. Scientific Reports. 2016;6.

35. Li H, Guo Q, Li Q, Yang L. Long-reads reveal that Rhododendron delavayi plastid genome contains extensive repeat sequences, and recombination exists among plastid genomes of photosynthetic Ericaceae. PeerJ. 2020.

36. Wang W, Yang T, Wang HL, Li ZJ, Ni JW, Su S, et al. Comparative and Phylogenetic Analyses of the Complete Chloroplast Genomes of Six Almond Species (Prunus spp. L.). Scientific Reports. 2020;10.

37. Palmer JD. Chloroplast DNA exists in two orientations. Nature. 1983;301.

38. Walker JF, Jansen RK, Zanis MJ, Emery NC. Sources of inversion variation in the small single copy (SSC) region of chloroplast genomes. American Journal of Botany. 2015;102.

39. Kim Y, Shin J, Oh DR, Kim AY, Choi C. Comparative analysis of complete chloroplast genome sequences and insertion-deletion (Indel) polymorphisms to distinguish five Vaccinium species. Forests. 2020;11.

40. Jakobsson M, Säll T, Lind-Halldén C, Halldén C. Evolution of chloroplast mononucleotide microsatellites in Arabidopsis thaliana. Theoretical and Applied Genetics. 2007;114.

41. George B, Bhatt BS, Awasthi M, George B, Singh AK. Comparative analysis of microsatellites in chloroplast genomes of lower and higher plants. Current Genetics. 2015;61.

42. Selkoe KA, Toonen RJ. Microsatellites for ecologists: A practical guide to using and evaluating microsatellite markers. Ecology Letters. 2006;9.

43. Nishiyama S, Fujikawa M, Yamane H, Shirasawa K, Babiker E, Tao R. Genomic insight into the developmental history of southern highbush blueberry populations. Heredity. 2021;126.

44. Kulkarni KP, Vorsa N, Natarajan P, Elavarthi S, Iorizzo M, Reddy UK, et al. Admixture analysis using genotyping-by-sequencing reveals genetic relatedness and parental lineage distribution in highbush blueberry genotypes and cross derivatives. International Journal of Molecular Sciences. 2021;22.

45. Bian Y, Ballington J, Raja A, Brouwer C, Reid R, Burke M, et al. Patterns of simple sequence repeats in cultivated blueberries (Vaccinium section Cyanococcus spp.) and their use in revealing genetic diversity and population structure. Molecular Breeding. 2014;34.

46. Brevis PA, Bassil N v., Ballington JR, Hancock JF. Impact of wide hybridization on highbush blueberry breeding. Journal of the American Society for Horticultural Science. 2008;133.

47. Coville FV. Improving the wild blueberry. In: G. Hambidge Editor. USDA yearbook of agriculture. Washington, U.S. Govt. Printing Office. 1937. p 559–574.

48. Sharpe RH. Horticultural development of Florida blueberries. Proceedings of the Florida State Horticultural Society. 1953;66.

49. Goldy RG, Lyrene PM. Meiotic abnormalities of Vaccinium ashei × Vaccinium darrowi hybrids. Canadian Journal of Genetics and Cytology. 1984;26.

50. Magdy M, Ou L, Yu H, Chen R, Zhou Y, Hassan H, et al. Pan-plastome approach empowers the assessment of genetic variation in cultivated Capsicum species. Horticulture Research. 2019;6.

51. Lyrene PM, Olmstead JW. The Use of Inter-Sectional Hybrids in Blueberry Breeding. International Journal of Fruit Science. 2012;12.

52. Rousi A. Hybridization between Vaccinium uliginosum and cultivated blueberry. Annales Agriculturae Fenniae. 1963;2.

53. Colle M, Leisner CP, Wai CM, Ou S, Bird KA, Wang J, et al. Haplotype-phased genome and evolution of phytonutrient pathways of tetraploid blueberry. GigaScience. 2019;8.

54. Chaisson MJ, Tesler G. Mapping single molecule sequencing reads using basic local alignment with successive refinement (BLASR): Application and theory. BMC Bioinformatics. 2012;13.

55. Quinlan AR, Hall IM. BEDTools: A flexible suite of utilities for comparing genomic features. Bioinformatics. 2010;26.

56. Koren S, Walenz BP, Berlin K, Miller JR, Bergman NH, Phillippy AM. Canu: Scalable and accurate long-read assembly via adaptive κ-mer weighting and repeat separation. Genome Research. 2017;27.

57. Hunt M, Silva N de, Otto TD, Parkhill J, Keane JA, Harris SR. Circlator: Automated circularization of genome assemblies using long sequencing reads. Genome Biology. 2015;16.

58. GCpp. https://github.com/PacificBiosciences/gcpp. Accessed 21 Feb 2022.

59. Walker BJ, Abeel T, Shea T, Priest M, Abouelliel A, Sakthikumar S, et al. Pilon: An Integrated Tool for Comprehensive Microbial Variant Detection and Genome Assembly Improvement. PLoS ONE. 2014;9.

60. Dierckxsens N, Mardulyn P, Smits G. NOVOPlasty: De novo assembly of organelle genomes from whole genome data. Nucleic Acids Research. 2017;45.

61. Marçais G, Delcher AL, Phillippy AM, Coston R, Salzberg SL, Zimin A. MUMmer4: A fast and versatile genome alignment system. PLoS Computational Biology. 2018;14.

62. Bankevich A, Nurk S, Antipov D, Gurevich AA, Dvorkin M, Kulikov AS, et al. SPAdes: A New Genome Assembly Algorithm and Its Applications to Single-Cell Sequencing. Journal of Computational Biology. 2012;19.

63. Huang X, Madan A. CAP3: A DNA sequence assembly program. Genome Research. 1999;9.

64. Tillich M, Lehwark P, Pellizzer T, Ulbricht-Jones ES, Fischer A, Bock R, et al. GeSeq - Versatile and accurate annotation of organelle genomes. Nucleic Acids Research. 2017;45.

65. Liu C, Shi L, Zhu Y, Chen H, Zhang J, Lin X, et al. CpGAVAS, an integrated web server for the annotation, visualization, analysis, and GenBank submission of completely sequenced chloroplast genome sequences. BMC Genomics. 2012;13.

66. Singh BP, Kumar A, Kaur H, Singh H, Nagpal AK. CpGDB : A Comprehensive Database of Chloroplast Genomes. Bioinformation. 2020;16.

67. Beier S, Thiel T, Münch T, Scholz U, Mascher M. MISA-web: A web server for microsatellite prediction. Bioinformatics. 2017;33.

68. Lohse M, Drechsel O, Kahlau S, Bock R. OrganellarGenomeDRAW--a suite of tools for generating physical maps of plastid and mitochondrial genomes and visualizing expression data sets. Nucleic acids research. 2013;41.

69. Frazer KA, Pachter L, Poliakov A, Rubin EM, Dubchak I. VISTA: Computational tools for comparative genomics. Nucleic Acids Research. 2004;32.

70. Rice P, Longden L, Bleasby A. EMBOSS: The European Molecular Biology Open Software Suite. Trends in Genetics. 2000;16.

71. Larkin MA, Blackshields G, Brown NP, Chenna R, Mcgettigan PA, McWilliam H, et al. Clustal W and Clustal X version 2.0. Bioinformatics. 2007;23.

72. NCBI Multiple Sequence Alignment Viewer. https://www.ncbi.nlm.nih.gov/projects/msaviewer/. Accessed: 21 Feb 2022.

73. Bi G, Mao Y, Xing Q, Cao M. HomBlocks: A multiple-alignment construction pipeline for organelle phylogenomics based on locally collinear block searching. Genomics. 2018;110.

74. Cognato AI, Vogler AP. Exploring data interaction and nucleotide alignment in a multiple gene analysis of Ips (Coleoptera: Scolytinae). Systematic Biology. 2001;50.

75. Minh BQ, Schmidt HA, Chernomor O, Schrempf D, Woodhams MD, von Haeseler A, et al. IQ-TREE 2: New Models and Efficient Methods for Phylogenetic Inference in the Genomic Era. Molecular Biology and Evolution. 2020;37.

76. Hoang DT, Chernomor O, von Haeseler A, Minh BQ, Vinh LS. UFBoot2: Improving the ultrafast bootstrap approximation. Molecular Biology and Evolution. 2018;35.

77. Letunic I, Bork P. Interactive tree of life (iTOL) v5: An online tool for phylogenetic tree display and annotation. Nucleic Acids Research. 2021;49.

78. Leigh JW, Bryant D. POPART: Full-feature software for haplotype network construction. Methods in Ecology and Evolution. 2015;6.

79. Clement M, Snell Q, Walke P, Posada D, Crandall K. TCS: Estimating gene genealogies. In: Proceedings - International Parallel and Distributed Processing Symposium, IPDPS 2002.

