## Additional File 1 for "Chloroplast genome assemblies and comparative analyses of major *Vaccinium* berry crops"

### Additional File 1 - Supplementary Figures

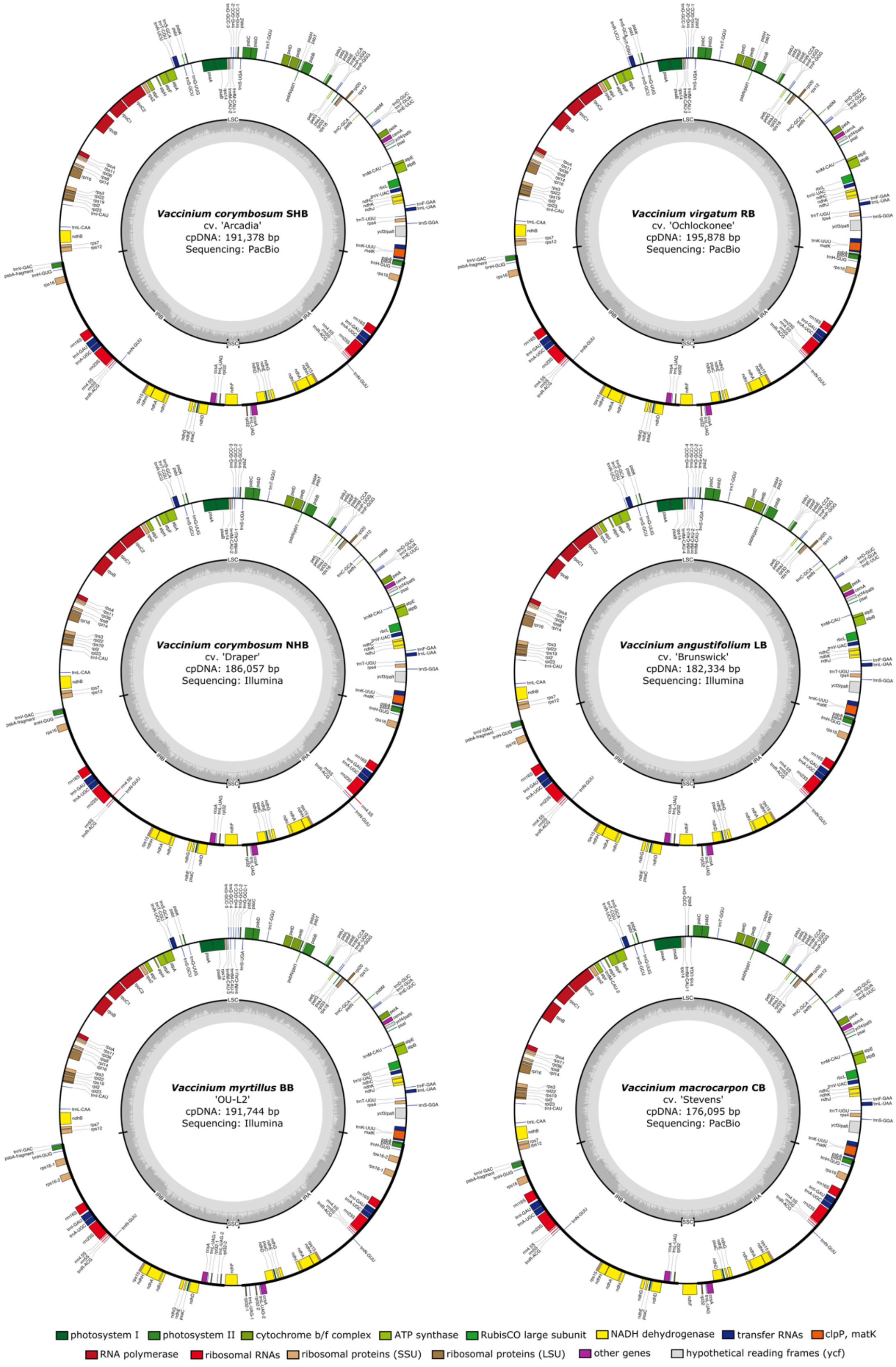

**Figure S1.** Circular chloroplast genome maps of six *Vaccinium* species. Five species were assembled de-novo and annotated here: southern highbush blueberry (SHB), rabbiteye blueberry (RB), northern highbush blueberry (NHB), lowbush blueberry (LB) and bilberry (BB). Cranberry (CB) was assembled previously (Diaz-Garcia et al., 2019) and reannotated here. Genes drawn outside and inside the maps represent genes transcribed counterclockwise and clockwise, respectively, and the different colors represent their functional annotation. The large single copy (LSC), inverted repeats (IRA and IRB), and small single copy (SSC) regions are shown in the black inner circle. The gray inner circle shows GC content.

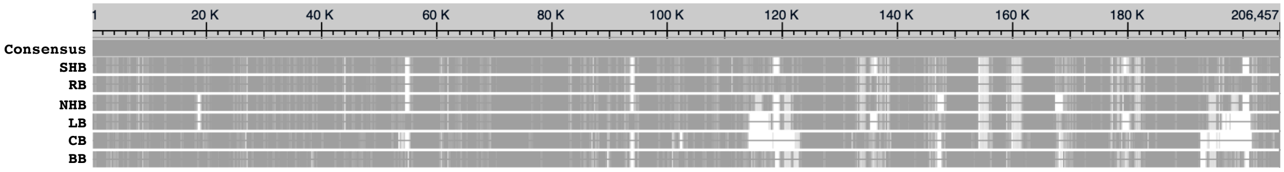


**Figure S2.** ClustalW multiple sequence alignment of the complete plastomes of six *Vaccinium* species. SHB: southern highbush blueberry, RB: rabbiteye, NHB: northern highbush blueberry, LB: lowbush blueberry, CB: cranberry, BB: bilberry. 'Consensus' refers to the consensus sequence obtained from the alignment.

```
SSR1 (TAC)
Southern highbush      TACTATTA-----G
Rabbiteye              TACTATTA-----G
Northern highbush      TACTATTA-----G
Lowbush                TACTATTA-----G
Bilberry               TACTACTACTACTA
Cranberry              TACTATTA-----G

SSR2 (TA)
Southern highbush      TATATATAT--ATA
Rabbiteye              TATATATAT--ATA
Northern highbush      TATATATAT--ATA
Lowbush                TATATATAT--ATA
Bilberry               TATATATATCTATA
Cranberry              TATATATATCTATA

SSR3 (AT)
Southern highbush      ATATATATATAT
Rabbiteye              ATATATATATAT
Northern highbush      ATATATATATAT
Lowbush                ATATATATATAT
Bilberry               AT------ATAT
Cranberry              AT------ATAT

SSR4 (AAG)
Southern highbush      AAGAAG------G
Rabbiteye              AAGAAG------G
Northern highbush      AAGAAG------G
Lowbush                AAGAAG------G
Bilberry               AAGAAG------G
Cranberry              AAGAAGAAGAAGG

SSR5 (TTTCTT)
Southern highbush      TTTCTTTTTCTTTTTCTTT------C
Rabbiteye              TTTCTTTTTCTTTTTCTTTTTCTTTC
Northern highbush      TTTCTTTTTCTTTTTCTTTTTCTTTC
Lowbush                TTTCTTTTTCTTTTTCTTTTTCTTTC
Bilberry               TTTCTTTTTCTTTTTCTTT------C
Cranberry              TTTCTTTTTCTTTTTCTTT------C

SSR6 (AT)
Southern highbush      AT-------------------------------------------ATATATAT
Rabbiteye              AT-------------------------------------------ATATATAT
Northern highbush      AT-------------------------------------------ATATATAT
Lowbush                AT-------------------------------------------ATATATAT
Bilberry               ATATATATACCATAATATATATATATATTATTTATATACCATGATATATATAT
Cranberry              ATGT-----------------------------------------ATATATAT

SSR7 (TA)
Southern highbush      TATATATATAAA
Rabbiteye              TATATATATAAA
Northern highbush      TATATATATAAA
Lowbush                TATATATATAAA
Bilberry               TATATATATATA
Cranberry              TCTATATATATA
```

**Figure S3** continues...

```
SSR8 (TA)
Southern highbush      TATATATATAAAA----------------------------T
Rabbiteye              TATATATATAAAA----------------------------T
Northern highbush      TATATATATAAAA----------------------------T
Lowbush                TATATATATAAAA----------------------------T
Bilberry               TATTTATATATAT------TAACTCAATTATCTCAGTCATTT
Cranberry              TATTTATATATATATATATTAACTTAATTATCTCAGTCATTT

SSR9 (TA)
Southern highbush      TATATAT----------ATATATAAT--A
Rabbiteye              TATATAT----------ATATATAAT--A
Northern highbush      TATATAT----------ATATATAAT--A
Lowbush                TATATAT----------ATATATAAT--A
Bilberry               TATATAT------------ATATAAT--A
Cranberry              TATATATTATTTATTATATATATAAATAA

SSR10 (TA)
Southern highbush      TATAT------ATAT--T
Rabbiteye              TATAT------ATAT--T
Northern highbush      TATAT------ATAT--T
Lowbush                TATAT------ATAT--T
Bilberry               TATATGTATATATATATT
Cranberry              TATAT------ATATATT

SSR11 (TAT)
Southern highbush      TTTAAAAAAGGATATTATTATTATTA
Rabbiteye              TTTAAAAAAGGATATTATTATTATTA
Northern highbush      TTTTAAAAAGGATATTATTATTATTA
Lowbush                TTTTAAAAAGGATATTATTATTATTA
Bilberry               TTTTAAAAAGGATATTATT------A
Cranberry              T------AAGTATATTATT------A
```

**Figure S3.** Sequence variability of 11 SSRs identified in six *Vaccinium* species. For each SSR, the underlined sequence shows the motif identified by the MISA software. SHB: southern highbush blueberry, RB: rabbiteye blueberry, NHB: northern highbush blueberry, LB: lowbush blueberry, CB: cranberry, BB: bilberry.
